# Endosome transcriptomics reveal trafficking of Cajal bodies into multivesicular bodies

**DOI:** 10.1101/2025.05.22.655499

**Authors:** Jasleen Singh, Justin Krish Williams, Quinn Elliott, Rohit Jhawar, Lucas Ferguson, Kathleen Collins, Randy Schekman

## Abstract

All eukaryotic cells secrete exosomes, a type of extracellular vesicles (EVs) derived from the endocytic compartments known as multivesicular bodies (MVBs), or late endosomes (LEs). Exosomes contain a diverse range of cargo such as nucleic acids, proteins, lipids and small molecules but whether these contents have a biological function remains an area of intense investigation. Over the last decade, numerous studies have described the transcriptome of exosomes but very little is known about the RNA content of the MVBs, the source compartment for exosome biogenesis. Here we determine the small-RNA transcriptome of highly purified MVBs and report that various classes of nuclear small regulatory RNAs such as small-Cajal body associated RNAs (scaRNAs), small-nucleolar RNAs (snoRNAs) and small-nuclear RNAs (snRNAs) traffic to MVBs. We show that this RNA-trafficking requires the function of ESCRT machinery but is independent of canonical LC3 lipidation mediated selective autophagy. Furthermore, blocking the activity of a PI3K Class 3 enzyme, VPS34, required for recruitment of the ESCRT machinery to the endosome, prevents the turnover of these nuclear RNAs in MVBs. Our results provide a mechanism for targeting nuclear ribonucleoprotein complexes (RNPs), such as Cajal bodies, for degradation and turnover by the cytoplasmic endo-lysosomal pathway.

**Significance Statement:** Endosomes are cytoplasmic, membrane-bound subcellular organelles that are sites for biogenesis of exosomes, a class of extracellular vesicles, thought to mediate intercellular communication via their packaged cargo such as RNA. Previous studies have focused on the transcriptome of exosomes however very little is known about the identity of RNAs and mechanisms by which they are sorted into endosomes. Here we report a comprehensive endosome transcriptome and provide evidence that several nuclear RNA-protein complexes (RNPs) sort into endosomes, a previously unappreciated phenomenon. We show that this process requires the activity of endosomal sorting complexes and phospholipids characteristic of cellular endocytic compartments. Our study provides a mechanism for recycling and disposal of unwanted nuclear RNPs by the cytoplasmic endolysosomal pathway.

## Introduction

All eukaryotic cells secrete diverse types of extracellular vesicles (EVs) derived from an equally varied set of cellular membranes, such as microvesicles that bud from the plasma membrane (PM) and exosomes, originating from endocytic compartments of the cell (1–4). Whereas the contents of microvesicles are largely representative of the general cytoplasm, exosomal cargo appears to be distinct from the overall cytoplasmic contents implying underlying active cellular processes that mediate selective cargo sorting into exosomes (5). Indeed, several RNA-binding proteins (RBPs) form a part of the cellular machinery aiding in the selective sorting of microRNAs (miRNAs) into exosomes (5–10). Exosomes along with their cargo released from one cell are routinely observed to be taken up by other cells giving rise to an idea that exosomes mediate cell-cell communication via transfer of functional biomolecules (11, 12). Of particular interest are small-RNAs (sRNA) such as miRNAs and piRNAs that have the potential to relay important gene regulatory information to the recipient cells (13, 14). This anticipation for the putative functions of exosomes has led to a myriad of studies exploring the RNA content of exosomes and other EVs but whether there is a direct causal relationship between a particular RNA cargo and the function it elicits in the recipient cells has not been established (2, 15). In contrast to exosomes, very little is known about the RNA content of endosomes, the compartment for biogenesis of exosomes, and its relationship to the RNA content in secreted exosomes.

Endosomes are membrane-bound organelles of the endocytic pathway that play a crucial role in sorting of membrane proteins, lipids and other biomolecules to their appropriate cellular destinations thus controlling fundamental cellular processes such as immune signaling, nutrient sensing and uptake, cell-adhesion and membrane recycling (16, 17)5/22/2025 11:31:00 AM. Depending on their morphology, composition and localization, endosomes are primarily characterized as early (EEs), recycling (REs) and late endosomes (LEs) (17–19). Generally proximal to the PM, EEs and REs aid in the transport of housekeeping membrane proteins back to the PM. EEs undergo a complex set of maturation steps giving rise to LEs containing unwanted membrane proteins and other biomolecules, at least a subset of which are targeted for degradation in lysosomes (17, 19). LEs and to a lesser extent EEs, contain intraluminal vesicles (ILVs) that are formed by inward budding of the endosomal limiting membrane and such endosomes containing ILVs are often termed as multivesicular bodies (MVBs) (20–23). ILV biogenesis is mediated by the action of endosomal sorting complexes required for transport (ESCRT; ESCRT 0, I, II and III) and several accessory proteins such as AAA-type ATPase VPS4 which together lead to the sorting of ubiquitylated cargo, endosome membrane remodeling and scission (24–30). MVBs and related compartments can also gain cargo via autophagy by fusing with autophagosomes to recycle unwanted cellular components in a process crucial for maintaining cellular homeostasis (31–33).

Interestingly, certain physiological triggers or experimental treatments of cells with drugs such as ionomycin (a calcium ionophore), bafilomycin A1 (inhibitor of lysosomal V-ATPase) and bacterial pore-forming toxins result in the MVB fusion with the PM instead of lysosomes, thus releasing the ILVs contained within MVBs into the extracellular environment (34–37). Such secreted ILVs are termed exosomes (2).

In addition to ubiquitylated proteins, several RNA molecules have been shown to selectively sort into ILVs in a process aided by specific RBPs and other unknown factors (5–8, 10). Notably, in such cases the RNA species of interest were initially identified in exosomes and subsequently shown to be selectively sorted into ILVs/MVBs (5, 6). Transcriptomes of exosomes prepared from various cell-types are reported routinely however, the overall RNA content of MVBs from any cell-type remains largely unknown. Given such an intricate connection between RNA content of exosomes and RNA sorting into MVBs, we sought to determine the transcriptome of both these compartments simultaneously.

Inspired by the previously published methods for organelle-immunoprecipitation (organelle-IP) (38–41), we developed a protocol for MVB-IP by engineering an affinity tag on CD63, a prominent MVB and exosome marker (5, 42). Next, we subjected the RNA isolated from purified MVBs to a highly sensitive method of sRNA-sequencing and determined that several nuclear non-coding RNAs including the components of Cajal bodies (CBs) such as small-Cajal body associated RNAs (scaRNAs) are sorted into MVBs. We provide evidence that such RNA sorting is associated with ILV budding into MVBs. Our study provides evidence that nuclear RNAs and RNPs such as CBs or their components are targeted for degradation and turnover by the cytoplasmic endo-lysosomal pathway, a previously unappreciated phenomenon.

## Results

### Purification and small-RNA sequencing of multivesicular bodies

To rapidly isolate MVBs, we generated a stable cell line expressing N-terminally 3xHA-tagged version of CD63, a routine MVB and exosome marker (Fig. 1A). MVBs from this cell line were immunoprecipitated using magnetic α-HA beads and subjected to immunoblot analysis, similar to previously described lysosome-IP protocols (Fig. 1A and Fig. S1A-C). The data shows a consistent enrichment of MVB membrane and lumen markers (Fig. 1A) and a depletion of markers for other organelles (Fig. S1A-C).

**Figure 1.**
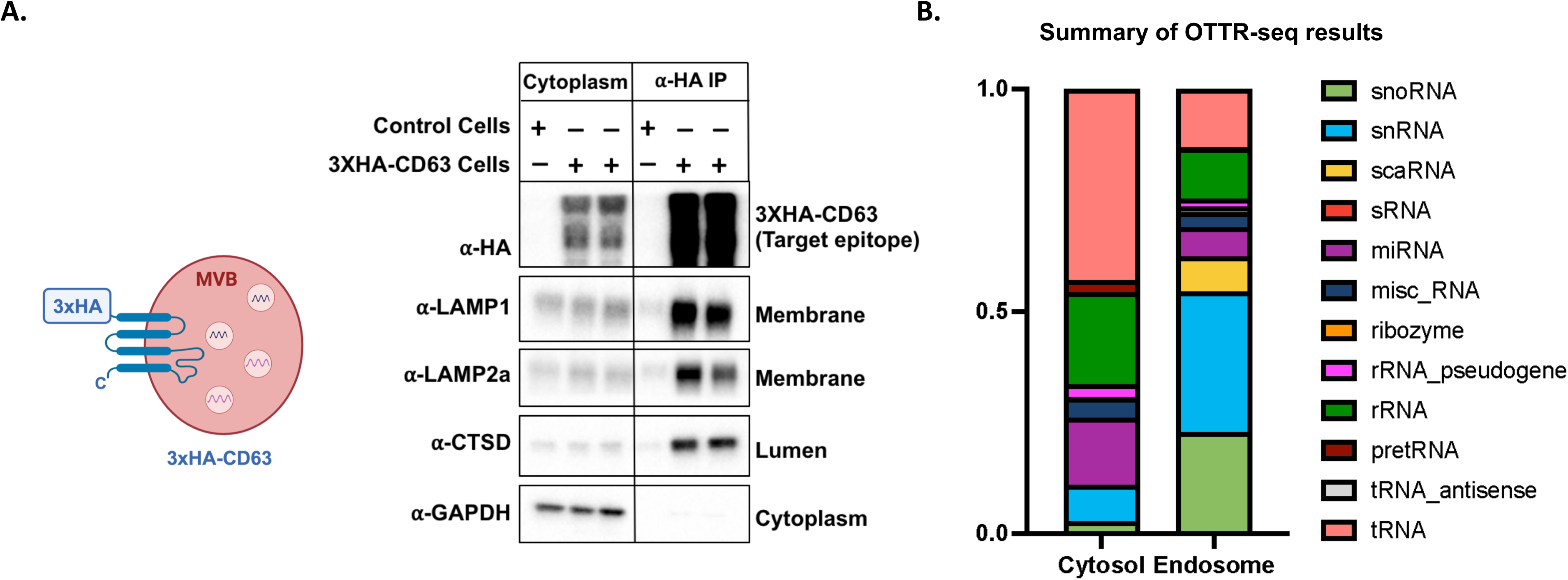
Purification and small-RNA sequencing of MVBs/LEs. A) Left, a cartoon showing the schematic of engineered MVBs/LEs containing a 3xHA tag on the cytosol-facing N-terminus of an MVB marker, CD63; right, immunoblots against the target epitope, cytoplasmic and MVB markers in an untagged control line (HEK 293T) vs an engineered stable cell-line expressing 3xHA-CD63 (HEK 293T background), a comparison between the cytoplasmic lysate and anti-HA IP is shown. B) A summary of the OTTR-seq results.

To determine the sRNA transcriptome of isolated MVBs, we performed sRNA-seq using ordered two template relay (OTTR-seq) (43). A summary of the OTTR-seq data shows a significant enrichment of several nuclear non-coding RNAs such as scaRNAs, snRNAs, snoRNAs (Fig. 1B). To rule out a general nuclear contamination, we tested our MVB-IP samples for nuclear markers, Lamin A/C and Histone H4. Our results indicate an absence of these markers in MVB-IPs (Fig. S1C). Additionally, we analyzed the levels of nuclear pre-tRNAs in our OTTR-seq data set and consistent with a lack of general nuclear contamination, we found the pre-tRNA levels to be significantly lower in MVBs relative to cytoplasm (Fig. S1D).

In order to study the relationship between RNA trafficking into MVBs and the RNA content of EVs, we compared the sRNA contents of MVBs, the high-speed pellet (HSP) obtained by differential centrifugation of conditioned medium (CM) and the EVs immunoprecipitated from CM using a CD63 antibody (CD63^+^EVs). We found that although a significant proportion of RNA in the HSP and CD63^+^EVs overlapped with the general cytosolic contents, they contained various RNAs distinct from each other and MVBs. This suggested a rather complex relationship between MVB RNA trafficking and RNA content of EVs (Fig. S2).

### Several non-coding RNAs are detected in MVBs

Using curated and publicly available transcriptomes, we compared the OTTR-seq reads obtained from MVBs and cytoplasm and found a diverse array of RNA including nuclear-body RNAs such as scaRNAs, snoRNAs and snRNAs to be enriched in MVBs (Fig. 2A). We further compared the MVB and cytoplasm samples against an miRbase dataset to assess differential miRNA enrichment in these compartments and found that several miRNAs were selectively sorted into MVBs (Fig. 2B).

**Figure 2.**
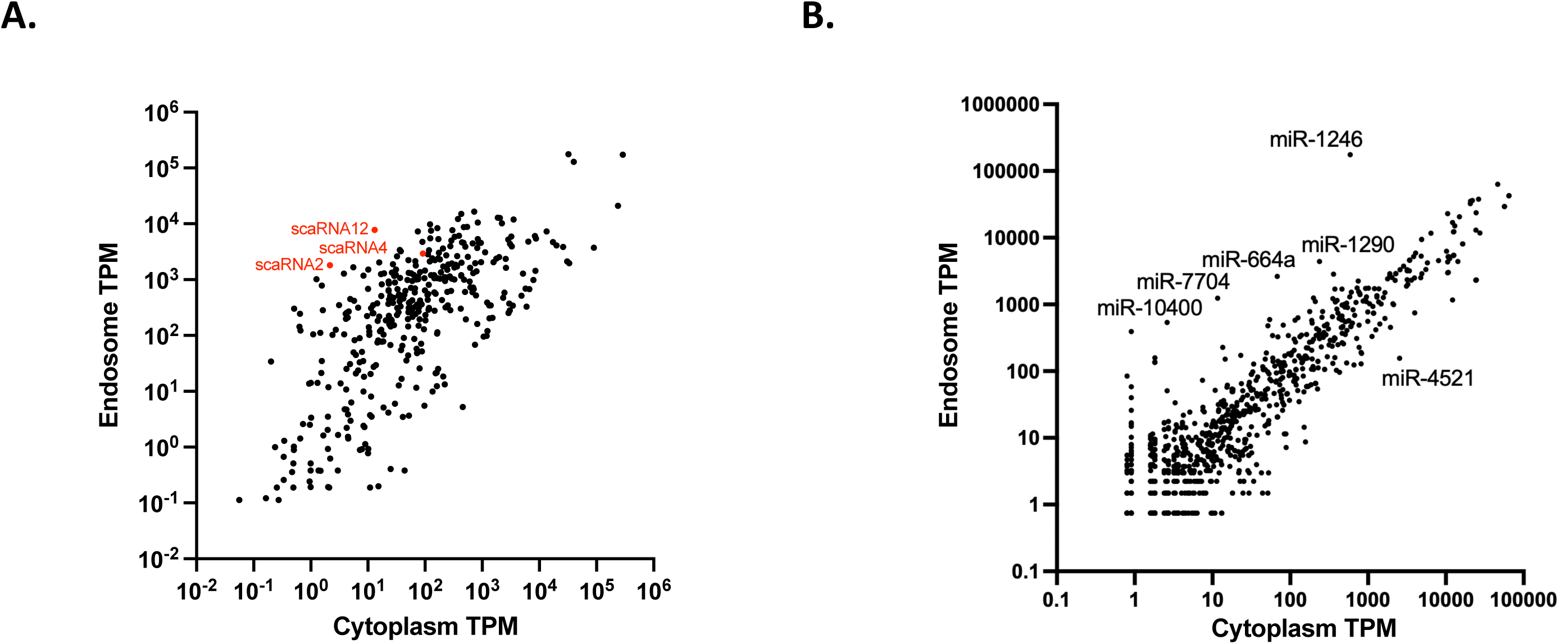
OTTR-seq of purified MVBs. A) A comparison of non-coding RNA transcriptome of purified MVBs and general cytoplasm. scaRNAs further analyzed in this study are highlighted. B) A comparison of miRNA content of purified MVBs and general cytoplasm.

To validate the presence of scaRNAs in the MVB-IPs, we developed independent reverse transcription PCR (RT-PCR) assays against scaRNAs 2, 4 and 12 (Fig. 3). The results show a modest enrichment of the full-length scaRNAs 2, 4 and 12 in the MVB-IPs relative to cytoplasm (Fig. 3A). This modest enrichment in RT-PCR assay is in contrast to high level of scaRNA enrichment observed using OTTR sRNA sequencing. Because OTTR-seq is exceptionally robust at detecting sRNA reads (43, 44), we reasoned that a high level of scaRNA enrichment in MVB-IPs could be a result of their processing into sRNA fragments by lysosomal ribonucleases, a feature missing in the cytoplasmic pool of scaRNAs. To test this idea, we subjected RNA prepared from MVB-IPs and cytoplasm samples to CORALL-seq, a technique capable of detecting longer transcripts, unlike OTTR-seq which preferentially captures smaller transcripts. Consistent with this idea, the level of full-length scaRNA enrichment detected by CORALL-seq in MVB-IPs relative to cytoplasm was greatly reduced compared to sRNA reads obtained by OTTR-seq (Fig. S4).

**Figure 3.**
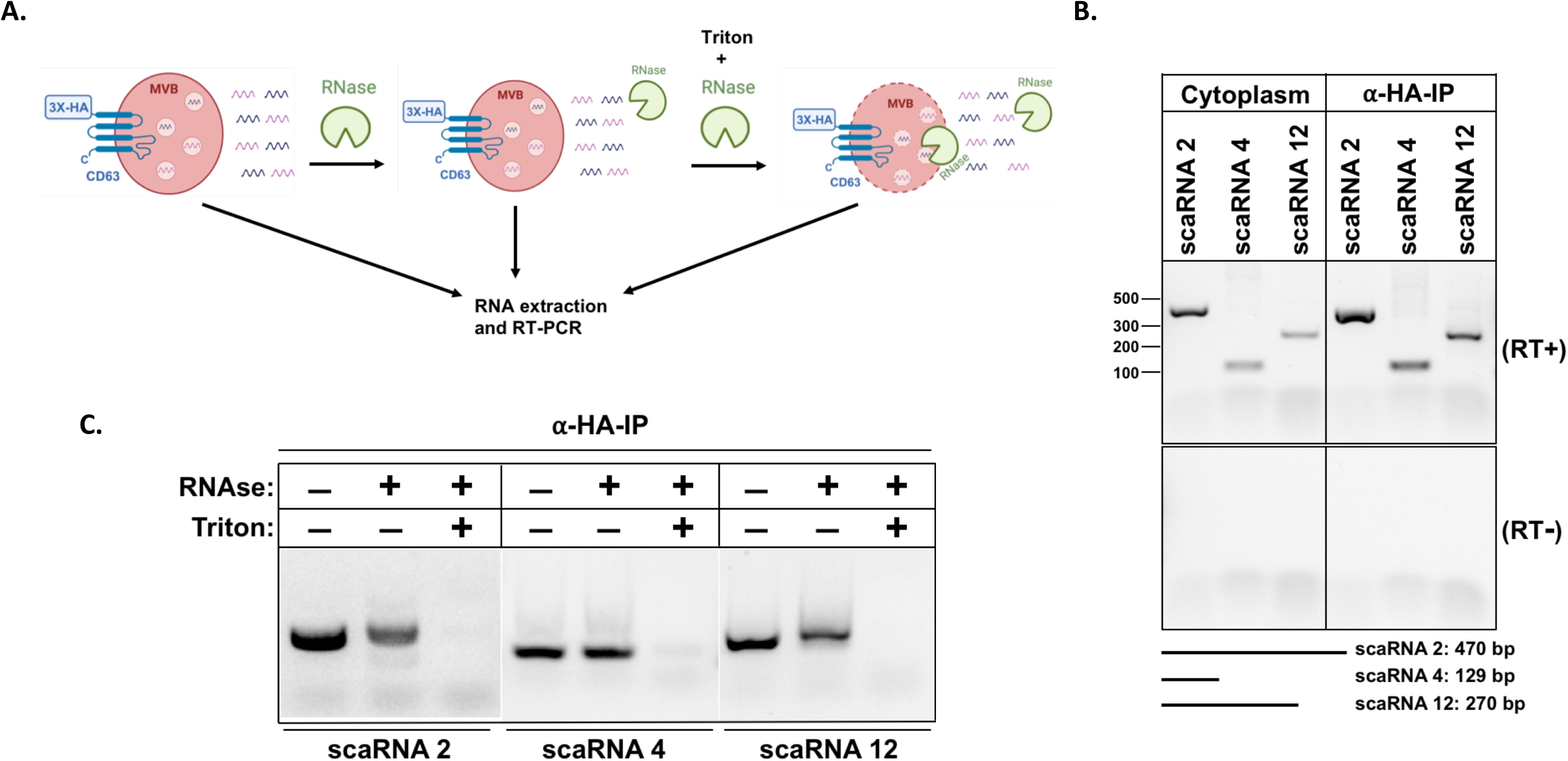
Small-Cajal body RNAs localize to MVBs. A) A schematic for the RNase treatments to test for the localization of scaRNAs relative to MVBs. B) RT-PCR analysis of scaRNAs 2, 4 and 12 in affinity purified MVBs relative to cytoplasm; no RT control (bottom panel); expected sizes of the scaRNAs are shown at the bottom of the gel. C) RNase treatments to test the localization of scaRNAs relative to MVBs, in presence or absence of Triton X-100.

### Small-Cajal body RNAs localize into MVBs

Our experiments suggested that scaRNAs traffic to MVBs but did not rule out the possibility of non-specific association of such RNAs with the outer surface of MVBs. To test the localization of scaRNAs within MVBs, we subjected our MVB-IP samples to RNase treatment in the presence or absence of the detergent Triton X-100 (Fig. 3A). scaRNAs localized within MVBs should be inaccessible to RNase treatment unless the membrane barrier is breached on addition of a detergent such as Triton X-100. The three RNAs tested in this experiment, scaRNAs 2, 4 and 12, were susceptible to RNase treatment only in the presence of detergent, consistent with their localization inside the MVBs (Fig. 3C).

### Bafilomycin A1 and ionomycin stimulate the secretion of scaRNAs in EVs

We previously showed that treatment of cells with a calcium ionophore, ionomycin, causes plasma membrane (PM) damage which mobilizes MVBs to fuse with and repair the damaged PM (35, 36). This process results in ILVs contained within MVBs to be released into the condition medium (CM) as exosomes. Similarly, it has been reported that bafilomycin A1 (BafA1) treatment of cells causes MVB and lysosomal deacidification resulting in the MVB fusion to PM and release of exosomes (34, 37). We reasoned that if scaRNAs are contained within the MVBs, treatment of cells with ionomycin and BafA1 should result in the secretion of these RNAs within exosomes detected in the CM (Fig. 4A). Indeed, treatment of cells with these drugs resulted in the secretion of scaRNAs 2 and 4 detected in an HSP fraction sedimented from the CM (Fig. 4A and B). To determine whether scaRNAs are contained inside the EVs, we subjected the HSP to RNase treatment in presence or absence of Triton X-100 (Fig. 4A and C). A significant proportion of scaRNAs in the HSP were insensitive to RNase treatment unless the HSP was treated with detergent, consistent with the scaRNA localization inside the secreted EVs (Fig. 4A and C).

**Figure 4.**
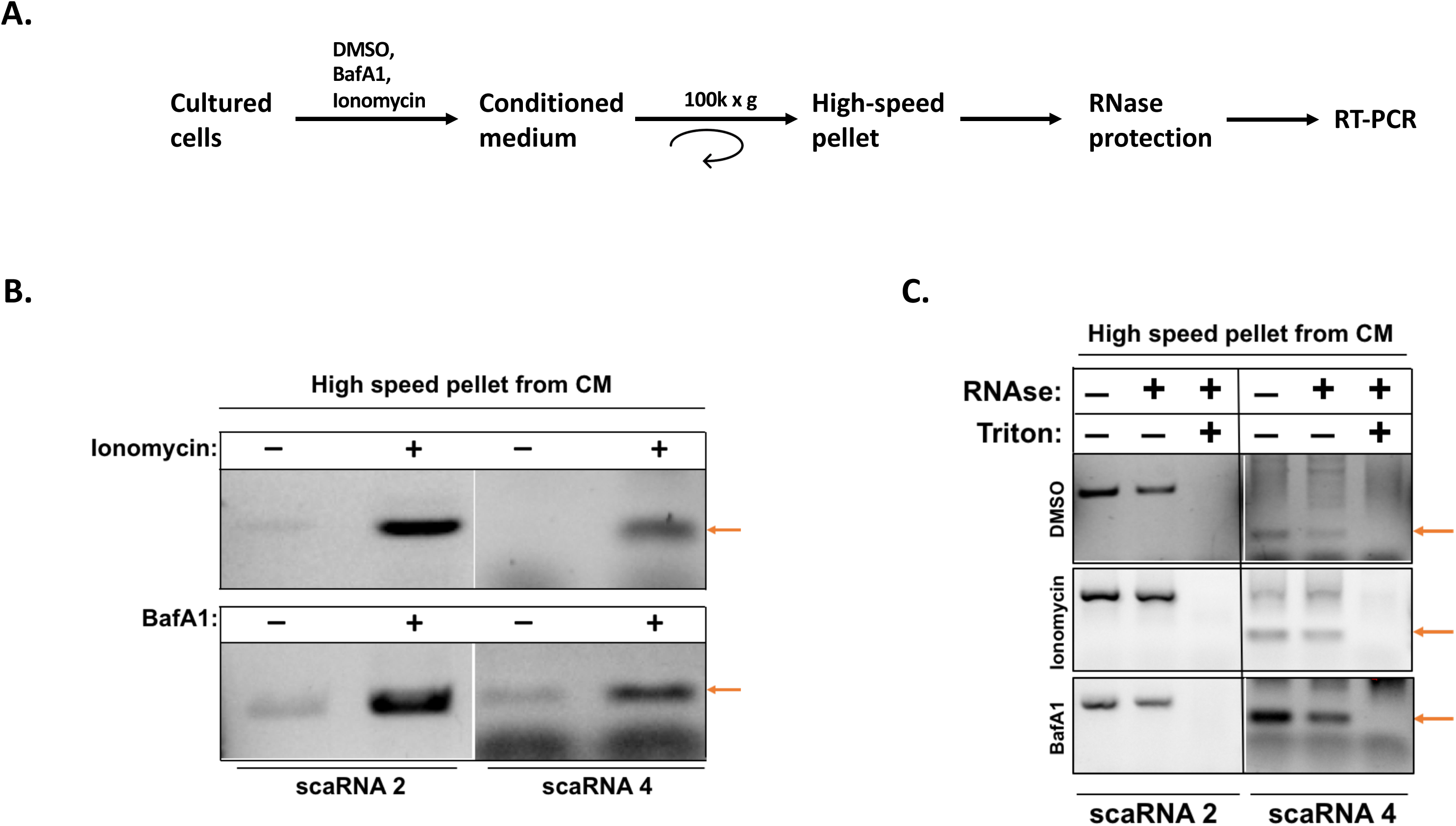
Ionomycin and bafilomycin A1 stimulate the secretion of scaRNAs in extracellular vesicles. A) A schematic for the experiments in B and C to test for the effect of ionomycin and bafA1 on scaRNAs secretion. B) Effect of ionomycin and bafA 1 on the accumulation of scaRNAs 2 and 4 in the extracellular fraction prepared by differential centrifugation of the conditioned medium. C) Test for the presence of scaRNAs 2 and 4 inside the EVs using RNase treatments in the presence or absence of Triton X-100. Red arrows in B and C indicate the expected positions of scaRNA 4.

### Coilin, a Cajal body marker, localizes to MVBs

We next examined the localization of a CB protein within MVBs. Immunofluorescence (IF) was used to visualize coilin, a key CB scaffolding protein (45–47), relative to CD63^+^ cytoplasmic compartments. Coilin (green puncta) co-localized with the CD63^+^ compartments (red circles) (Fig. 5 A and B and Supplement video 1). Additionally, coilin co-IP’d with MVBs, consistent with its localization to MVBs (Fig. 5C).

**Figure 5.**
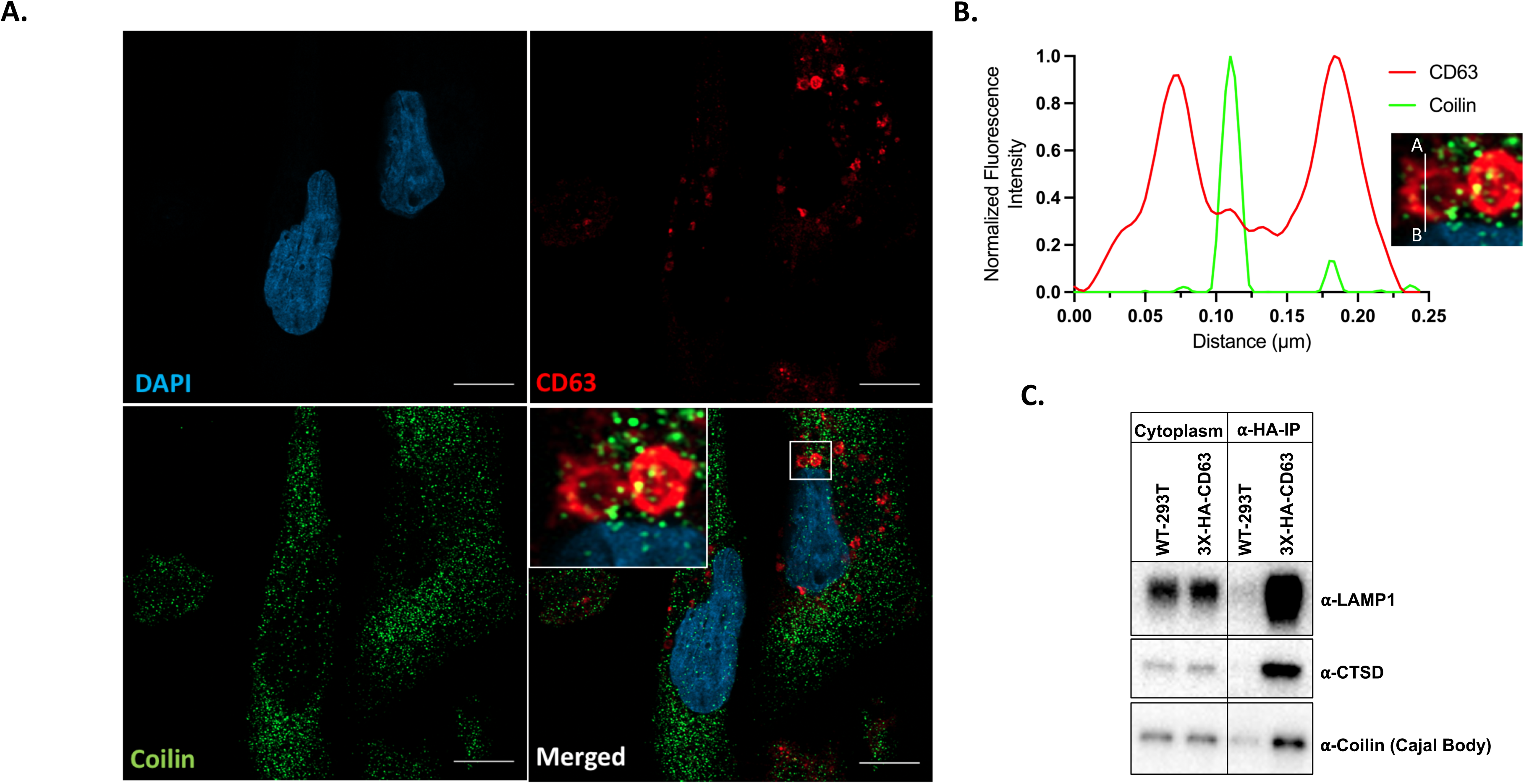
Coilin localizes to MVBs. A) Immunofluorescence assay showing the colocalization of coilin (green) to the CD63^+^ compartments (red). DAPI is shown in blue. B) Quantification of CD63 and coilin intensity from point A to point B in the inset of the figure 5A. C) Immunoblots to test for the co-immunoprecipitation of coilin with CD63^+^ positive MVBs.

### scaRNA localization to MVBs is independent of LC3 lipidation

Autophagy mediated lysosomal degradation of various biomolecules is a vital process required for cellular homeostasis. ATG7 is a key mediator of autophagy which is required for LC3 lipidation, a critical step in selective autophagy (31, 33). To test the possibility of autophagy mediated scaRNA localization to MVBs, we created a stable cell line expressing 3xHA-CD63 in an *atg7-/-* background and assayed for the levels of scaRNAs in MVB-IPs (Fig. S6). Our results show a clear block of LC3 lipidation in the 3xHA-CD63-*atg7-/-* whereas the scaRNA levels remained unchanged in the MVB IPs (Fig. S6 A and B). In an independent test, we induced autophagy by treating the cells with torin 1, an inhibitor of the mTOR kinase (48). The results showed that mTOR inhibition had no effect on the levels of scaRNAs in MVB-IPs, both in the wild-type and *atg7-/-* cells (Fig. S6 C and D).

### VPS4 is required for scaRNA sorting into MVBs

ESCRT complexes are essential for various steps of ILV biogenesis. ESCRT complexes aid in the sorting of cargo and promote inward budding into endosomes followed by VPS4 mediated scission of membrane buds to form ILVs (30). To test whether scaRNA sorting into MVBs is concomitant with ILV biogenesis, we transiently expressed a dominant negative allele of VPS4 mCherry (VPS4-DN) in a 3xHA-CD63 cell line and assessed the levels of scaRNAs in MVB-IPs. Our results showed that the expression of VPS4-DN did not interfere with the IP of MVBs (Fig. 6A) whereas, expression of the mutant significantly reduced the levels of scaRNAs 2 and 4 in MVB-IPs (Fig. 6 B and C).

**Figure 6.**
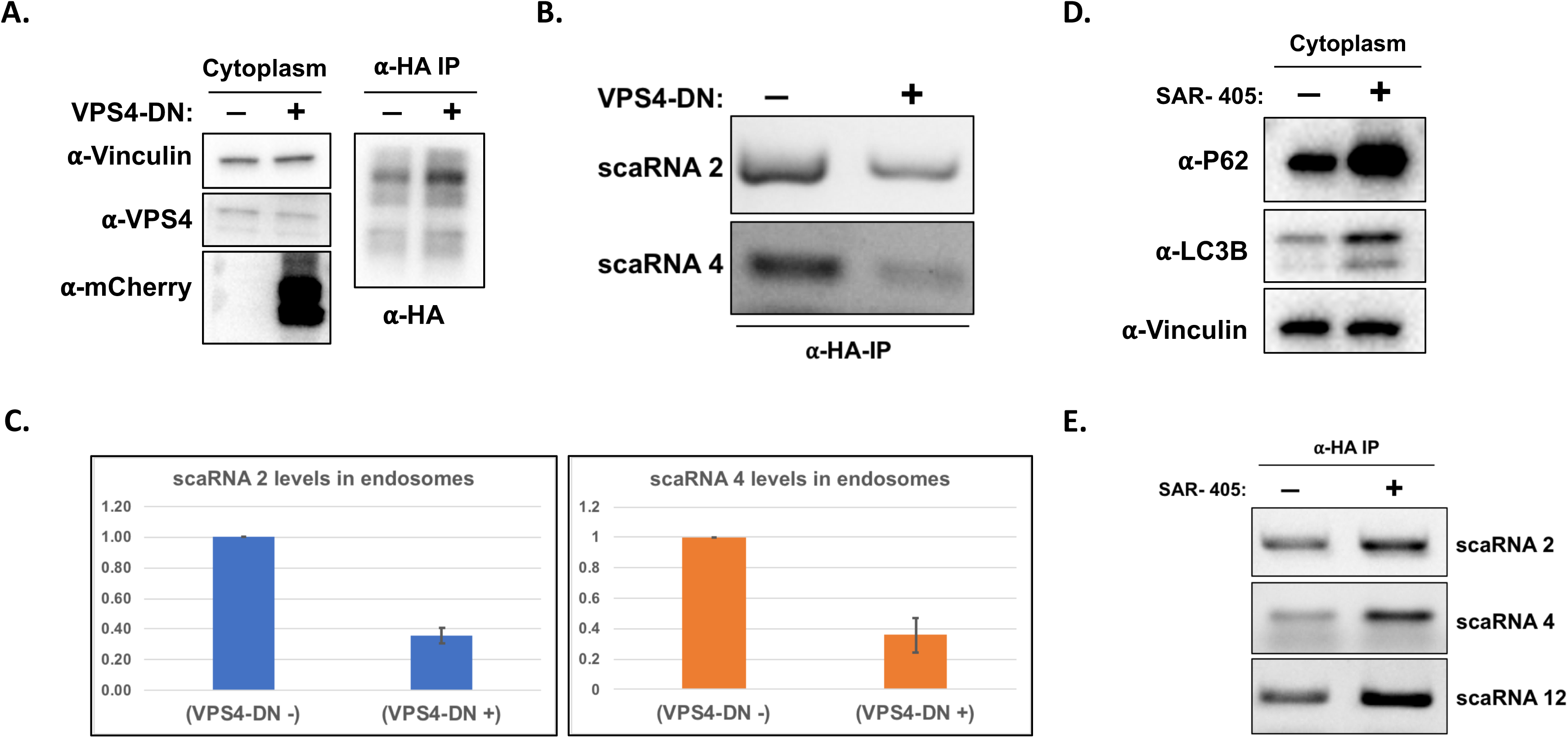
VPS4 and PI3P are required for MVB recruitment and turnover of scaRNAs, respectively. A) Immunoblots for the expression of an mCherry-VPS4-DN allele; native VPS4 and mCherry-VPS4-DN from cytoplasm (left) and purified MVBs (α-HA IP, right) are presented. B) RT-PCR assay to test for the effect of VPR4-DN expression on accumulation of scaRNAs in affinity purified MVBs. C) Quantification for the experiment in B. D) Immunoblots showing the effect of SAR-405, a VPS34 inhibitor, on the levels of P62 and LC3B. E) RT-PCR analysis to test for the levels of scaRNAs in MVBs upon treatment with SAR-405.

### Activity of PI3K, VPS34, is required for scaRNA turnover via endo-lysosomal pathway

Phosphatidylinositol-3-phosphate (PI3P) is a characteristic lipid of endo-lysosomal compartments of cells, essential for diverse aspects of lysosomal function (49). To test whether PI3P was required for endo-lysosomal turnover of scaRNAs, we treated 3xHA-CD63 cells with SAR-405, an inhibitor of PI3-kinase VPS34 (50). The treatment showed an expected increase in levels of a classic autophagy receptor, p62 (Fig. 6D). Importantly, SAR-405 treatment resulted in increased levels of scaRNAs 2, 4 and 12 in MVB-IPs consistent with the requirement of PI3P for efficient turnover of scaRNAs in endo-lysosomal compartments (Fig. 6E).

## Discussion

The RNA cargo contained within exosomes has been proposed to mediate aspects of cell-cell communication by eliciting gene regulatory responses in the recipient cells or through a cell autonomous homeostatic mechanism designed to rid cells of unwanted RNAs and other cargo (2). Both mechanisms could aid normal physiological functions of cells such as growth, differentiation, adhesion, immune signaling, nutrient sensing, and regulate progression of certain pathological conditions including neurodegenerative diseases and various forms of cancers. However, the evidence for a direct relationship between exosomal RNA cargo and the proposed associated function is not convincing due in part to the methodologies involved in preparing exosomes and the choice of RNA molecules putatively ascribed to a particular function (2). Such choice of RNA depends on several factors, for instance, a particular RNA is likely to be enriched in exosomes relative to its overall cellular level governed by its selective sorting into MVBs. Exosomal RNA content and enrichment has been measured by transcriptomics of sedimentable fractions prepared from CM of diverse cell types and varying degrees of purity. Comparable analysis of the RNA content of isolated MVBs/endosomes has not yet been reported.

We addressed the issue of selective RNA sorting and report the transcriptome of highly purified MVBs. Notably, our MVB-IP protocol is rapid (∼30 min) and likely to preserve sensitive endosomal content such as RNA, relative to traditional multi-step isolation methods. Moreover, the MVB-IP protocol is modular and allows for the preparation of fairly homogenous pools of MVBs from multiple samples using independently tagged endosomal membrane markers, which in our case is tetraspanin CD63. Our results show several similarities and some differences between the RNA content of CD63^+^ MVBs and CD63^+^ EVs. Such differences may suggest a dynamic and active regulation of pools of MVBs that are either targeted for endo-lysosomal degradation and are thus retained in the cell and others that discharge at the PM resulting in exosome secretion.

Our MVB transcriptomics data uncovered several nuclear non-coding RNAs including scaRNAs, snRNAs and snoRNAs that sort into MVBs. These RNAs form a part of diverse multimolecular RNPs that guide processing and modifications of other RNAs: snoRNAs are known to localize to nucleoli and are required for ribosomal RNA maturation, snRNAs which associate with RBPs to form snRNPs that are important for mRNA splicing and scaRNAs, which resemble snoRNAs in structure and function and assemble into large multimeric structures called CBs that guide important modifications of snRNAs required for maturation of snRNPs and rRNAs (47, 51–55). The presence of nuclear RNAs in MVBs was unexpected, hence we sought to validate some of these targets by developing gene-specific RT-PCR assays. We show that several scaRNAs co-IP with and localize inside of MVBs based on our test for sensitivity to RNase treatments (Fig. 3 B and C). Notably, the enrichment of scaRNAs inside MVBs measured by RT-PCR assays and total-RNA-seq (Fig. 3B and Fig. S4) is diminished as compared to that observed using sRNA-seq (Fig. 1C and Fig. 2A). We suggest that scaRNAs when sorted into MVBs likely undergo a rapid turnover thus sRNA-seq may enrich for processed and truncated fragments of scaRNAs (Fig. S4). In addition, or alternatively, the increased scaRNA enrichment in sRNA seq using OTTR-seq may result from the use of RNase treatment of MVB samples but not of cytoplasmic RNA samples prior to the preparation of RNA libraries. Moreover, a majority of cytosolic scaRNA pool has not undergone processing by the endo-lysosomal pathway and is hence, less likely to be represented in sRNA-seq (materials and methods). In contrast, the reverse is true for conventional total-RNA-seq rendering a higher probability of unprocessed cytoplasmic scaRNAs to be included in the library. Smaller truncated versions of scaRNAs, as expected in the RNase treated MVBs, are less appropriate samples for evaluation by total-RNA-seq. Nonetheless, our observation that cellular treatments that enhance the secretion of exosomes also increase the secretion of scaRNAs in membrane protected vesicles (Fig. 4), lead us to conclude that scaRNAs are sorted into MVBs and turned over by the endo-lysosomal pathway (Fig. 7).

**Fig. 7.**
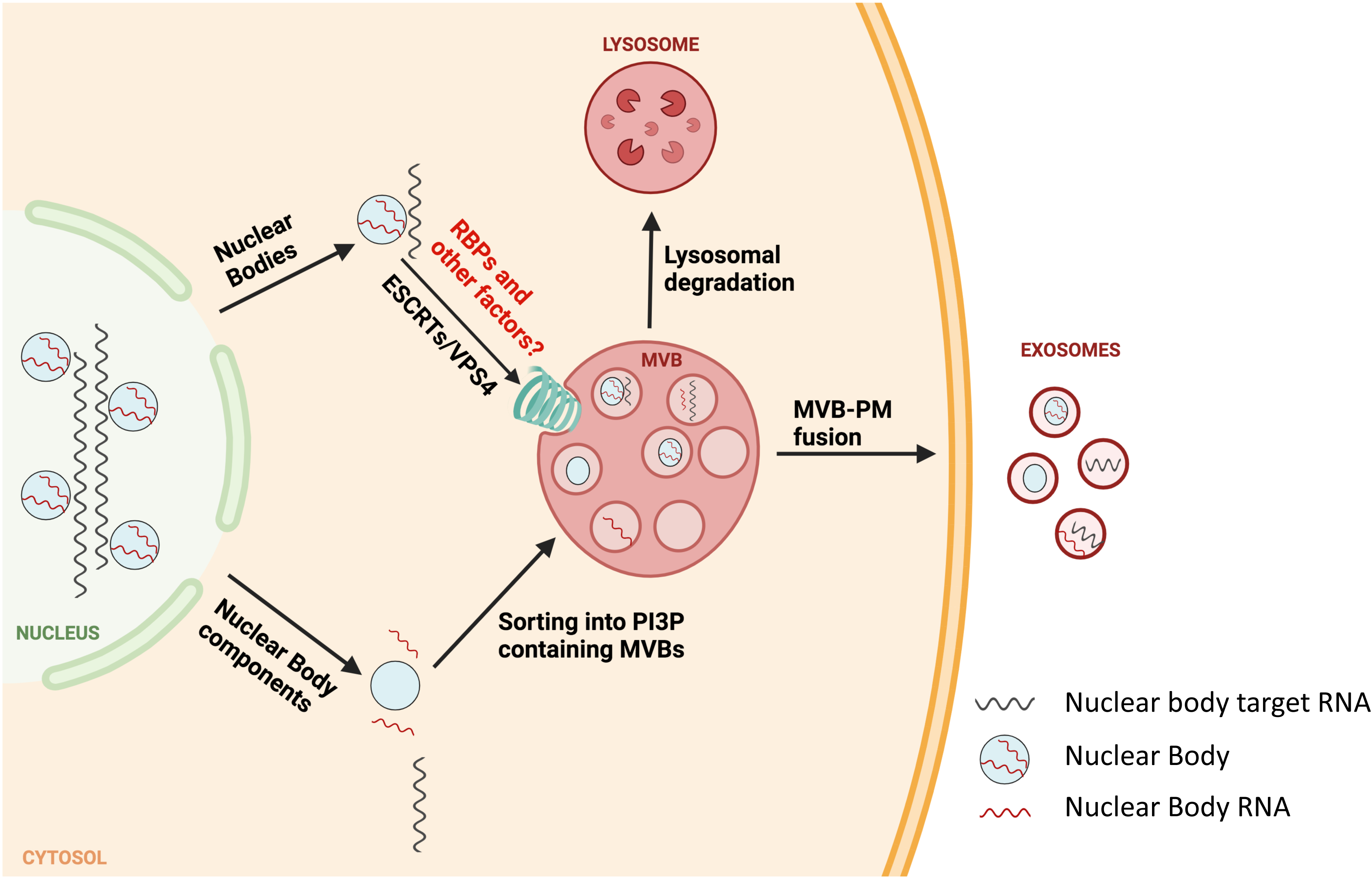
A model for recycling of nuclear body components in the MVBs.

CBs are primarily nuclear membraneless sub-organelles that aid in the maturation of snRNPs required for mRNA splicing (55). However, CB components such as coilin, a major structural component of CBs, and accessory proteins such as survival motor neuron (SMN 1), have previously been detected in the cytoplasm and likely undergo nucleus-cytoplasm shuttling (56–58). Additionally, CBs are known to disassemble during mitosis and as the nuclear envelope disintegrates during cell-division, CB components may mix with the surrounding cytoplasm (59, 60). Therefore, a subset of these components may be subject to turnover and the MVB sorting and endo-lysosomal turnover could aid this process. Consistent with this idea, we show that coilin localizes to and can be co-isolated with CD63^+^ MVBs (Fig. 5). Interestingly, several nucleolar RNPs and RNAs are known to associate with the perichromosomal layer (PCL) of mitotic chromosomes which aids in the nuclear compartmentalization of PCL associated RNPs and RNAs during the nuclear envelope reformation (61–63). However, it is possible that some nuclear RNPs and RNAs are inefficiently captured by this process and remain in the cytoplasm. In such a scenario, MVB sorting could play a role in the recycling of such nuclear components via endo-lysosomal pathway.

Coilin, along with other CB components, is also known to be a part of a related nuclear body called the histone locus body (HLB) that forms near all major histone gene loci where it guides the maturation of histone mRNAs (64–66). Notably, histone mRNAs are some of the most highly represented transcripts in our MVB total-RNA-seq (Fig. S5). These observations together with our discovery of snRNAs and snoRNAs in MVBs suggests that MVB sorting and endo-lysosomal turnover may play a broad role in maintenance of nuclear body homeostasis (Fig. 7). Autophagy related pathways are an important mechanism for recycling of unwanted cellular components including RNA. Autophagy is involved in recycling bulk cellular components in response to certain cues such as nutrient starvation but also may be targeted to selective targets (33). Selective autophagy of rRNA has been previously observed in *C. elegans* and yeast and is thought to maintain nucleotide homeostasis during development (67–70). Recently, an LC3-dependent EV loading and secretion pathway (LDELS) was shown to be required for MVB sorting and subsequent EV secretion of several RBPs and their associated small non-coding RNAs (71). In contrast, our data suggests that the MVB sorting of representative nuclear body components is largely independent of autophagy (Fig. S6). We show that despite being independent of autophagy, MVB sorting of nuclear body components requires a functioning ESCRT machinery as evidenced by a reduction in MVB scaRNA levels upon expression of VPS4-DN, compromising its function in ESCRT-dependent ILV budding (Fig. 6). How the autophagy and MVB pathways of turnover capture distinct cargo molecules remains an open question.

## Materials and methods

### Cell-lines and cell-culture

HEK293T and MDA-MB-231 cells used in this study were cultured and maintained in Dulbecco’s Modified Eagle’s Medium (DMEM, Thermo Fisher Scientific, Waltham, MA, USA) supplemented with 10% fetal bovine serum (FBS) at 37°C with 5% CO_2_. HEK293T-*atg7-/-* were a gift from the lab of Jayanta Debnath (Department of Pathology and Laboratory Medicine, UCSF). For EV preparation, cells were cultured in DMEM containing EV depleted FBS.

### Cloning, transient expression, lentivirus production and stable cell-line generation

CD63 cDNA with an N-terminal 3xHA tag and Kozak sequence (3xHA-CD63) was cloned into a pLJM1 lentiviral vector at Age1 and EcoR1 sites with expression cassette under a CMV promoter. HEK293T cells were grown to about 50% confluency in a 6-well plate, transfected with 1.5 μg of pLJM1-3xHA-CD63 and 200 ng and 1.4 μg of packaging vectors pMD2.G and psPAX2, respectively using TransIT-LT1 transfection reagent (Mirus Bio) following manufacturer’s recommendations. Lentivirus was harvested 72 h post-transfection, filtered using a 0.45 μM syringe filter followed by snap freezing and storing as small aliquots at -80°C. Stable cell-lines, HEK293T-3xHA-CD63 and HEK293T-*atg7-/-*-3xHA-CD63 were generated by transducing HEK293T and HEK293T-*atg7-/-*, respectively, using the filtered lentivirus from above in presence of 8 μg/ml polybrene for 24 h at which point fresh complete medium was added. Stable transformants were selected using 1 μg/ml puromycin and verified by immunoblotting. For transient expression of VPS4-DN, 1μg of pLIX-403 expressing VPS4-DN was transfected into the HEK293T-3xHA-CD63 grown to 80% confluency on 100 mm culture dishes, using TransIT-LT1 transfection reagent (Mirus Bio) following manufacturer’s recommendations. pLIX-403-VPS4-DN was a gift from the lab of James Hurley (Department of Molecular and Cell Biology, UC Berkeley).

### Multivesicular body (MVB) isolation

MVBs were isolated using previously described lysosome and organelle IP protocols with some modifications. Briefly, HEK293T cells expressing 3xHA-CD63 were cultured to 90% confluency in 150 mm culture dishes (all the subsequent steps performed on ice using ice-cold buffers), washed twice with 10 ml of PBS, harvested in 5 ml MVB-IP buffer (136 mM KCl, 10 mM KH_2_PO_4_, pH 7.4) containing protease inhibitors (1mM 4-aminobenzamidine dihydrochloride, 1 µg/ml antipain dihydrochloride, 1 µg/ml aprotinin, 1 µg/ml leupeptin, 1 µg/ml chymostatin, 1 mM phenymethylsulfonly fluoride, 50 µM N-tosyl-L-phenylalanine chloromethyl ketone and 1 µg/ml pepstatin); TCEP 0.5mM and 6 mL of OptiPrep (Sigma) for every 100 ml of buffer. Cells were centrifuged at 700xg for 5 min and resuspended in 1 ml of MVB-IP buffer. Mechanical lysis of cells was performed by 5-10 passes of the resuspended cells through a 22G needle. Cell lysates were centrifuged at 1500xg for 5 min and a post-nuclear supernatant (PNS) was collected and small aliquots were saved for immunoblotting. The remainder supernatant was centrifuged at 6000xg for 6 min to sediment any mitochondria and other large organelles and the remaining supernatant was collected and incubated with 100 μl of α-HA magnetic beads (Pierce, Thermo Scientific) pre-equilibrated with MVB-IP buffer. The mixture was incubated on a rotating mixer for 10 min, beads were collected using a magnetic rack and were washed three times with MVB-IP buffer. The MVBs immobilized on beads were routinely assayed for quality using immunoblotting and were processed for RNA extraction.

### MVB RNA sequencing

Immunoisolated MVBs, HSP, CD63^+^ EVs and cytoplasm were all dissolved in Tri Reagent (Zymo Research) and RNA was extracted using DirectZol micro prep kits (Zymo Research) as per the manufacturer’s protocol including an on-column DNase digestion step. MVBs and CD63^+^ EVs were additionally subjected to RNase treatment with 2 units of RNase ONE (Promega) for 15 min at room temperature prior to RNA extraction. RNA was quantified using a bioanalyzer in the QB3 genomics core facility at University of California, Berkeley. Equimolar amounts of RNA were used to prepare small-RNA libraries using ordered two-template relay sequencing (OTTR-seq) as previously described (43, 44). Conventional total RNA-seq libraries were prepared using CORALL-seq total RNA seq with ribo-depletion (Lexogen) according to manufacturer’s protocol. The libraries were analyzed for quality and sequenced using Illumina NovaSeq 6000 instrument (Azenta) and a minimum of 25 million 150x paired end reads were obtained for each sample.

### Reverse transcription-polymerase chain reaction assay (RT-PCR assay)

Purified RNA from indicated sources in each experiment was subjected to gene-specific reverse transcription using Revertaid kit (Thermo Scientific) as per the manufacturer’s protocol to synthesize cDNA. cDNA was then subjected to PCR using Phusion master mix (Thermo Scientific) up to 18 cycles for MVB and cytoplasm fractions and up to 22 cycles for the HSP obtained by differential centrifugation of conditioned medium. The products were analyzed by standard agarose gel electrophoresis. Primers used for RT-PCR assays are detailed below. scaRNA2: gene-specific RT primer and reverse primer for PCR-CCAGATCAGAATCGCCTCGAT; forward PCR primer-GTTTTAGGGAGGGAGAGCGG scaRNA4: gene-specific RT primer and reverse primer for PCR-GAGTGTTGGGTAGTACAGTCAG; forward PCR primer-ACTGGAGGACTAAGAAGGCT scaRNA12: gene-specific RT primer and reverse primer for PCR-TGTCAGATCCAAGGTTGCGC; forward PCR primer-CAGGCTGATGAGACTAAGGCGA

### Test for sensitivity to RNase

Purified MVBs or HSP pellet fractions were treated with 2 units of RNase ONE (Promega) for 10 min at room temperature. Where indicated, Triton X-100 (Sigma) was added to a final concentration of 1% before the addition of RNase ONE. The reaction was stopped by addition of SDS to a final concentration of 0.2% w/v (Thermo Scientific) and Tri Reagent (Zymo Research). The RNA was extracted as above and subjected to analysis either by sequencing or RT-PCR.

### Bafilomycin A1 and ionomycin treatments

WT HEK293T or 3xHA-CD63 HEK293T cells were grown to 90% confluency on 150 mm culture dishes, washed with PBS and treated with either DMSO (vehicle) or 1 μM ionomycin (Cayman Chemical) dissolved in 15 ml of serum-free media for 20 min. For BafA1 experiments, washed cells were treated with DMSO (vehicle) or BafA1 (100 nM final concentration) dissolved in 15 ml of complete medium (EV depleted) and the treatment was carried out for 4 h. The CM from ionomycin and BafA1 treated cells were collected and HSP fractions were obtained by differential centrifugation as described below (EV isolation). HSPs were resuspended in 300 µl of MVB-IP buffer and equally distributed into three reactions for testing their sensitivity to RNase treatments as indicated. RNase treatment was carried out as above and scaRNAs were evaluated by RT-PCR. Additionally, for Fig. S5 A and B, MVBs were isolated from BafA1 treated 3xHA-CD63 HEK293T cells and analyzed by RT-PCR

### Autophagy inhibition by torin

3xHA-CD63 HEK293T cells grown in 150 mm dishes to 90% confluency were treated with 10 nM torin (Cell Signaling Technologies) for 1 h. Treatments were evaluated by the inhibition of S6K phosphorylation relative to total S6K levels using immunoblotting. After treatment, MVBs were prepared as described above and assayed for scaRNA levels using RT-PCR.

### PI3 kinase, VPS34, inhibition by SAR-405

3xHA-CD63 HEK293T cells were grown in 150 mm dishes to a confluency of 90% and treated with SAR-405 (10 μM final concentration), an inhibitor of VPS34, for 3 h. Treatment was evaluated by examining the levels of P62 and LC3 using immunoblotting. MVBs were isolated post-treatment and assayed for scaRNA levels by RT-PCR as described above.

### Immunoblotting

Immunoblotting was performed according to routine procedures. Briefly, samples dissolved in Laemmli buffer (1x final concentration) were resolved by SDS-PAGE using 4-20% gradient Tris-glycine gels (Life Technologies). Proteins were transferred to polyvinylidene difluoride membranes (PVDF; EMD Millipore), blocked with 5% w/v non-fat dry milk powder in TBS-T at room temperature. Blocked membrane was washed three times with TBST and incubated overnight at 4°C with primary antibodies dissolved in 5% w/v bovine serum albumin (BSA) in TBS-T. Membranes were washed three times with TBS-T and incubated with secondary rabbit or mouse antibodies conjugated with horse radish peroxidase (HRP) at 1:10000 dilution in 5% milk TBS-T and then washed again three times with TBS-T. Signals were developed using ECL-2 or ECL picoPLUS HRP substrates as per manufacturer’s guidelines and blots were imaged using Biorad ChemiDoc^TM^ system.

All the primary antibodies used in this study are listed below and were at 1:1000 dilution in 5% BSA-TBST unless otherwise stated. Mouse α-HA (Cell-signaling technologies/CST C29F4), rabbit α-LAMP1 (CST D2D11), rabbit α-LAMP2a (Abcam 18528), rabbit α-Cathepsin D (CST 69854), rabbit α-Coilin (CST D2L3J), rabbit α-Vinculin (Abcam ab129002), rabbit α-VPS4 (Proteintech 14272-1-AP), rabbit α-mCherry (Novus NBP2-25157), mouse α-P62/SQSTM1 (Abcam ab56416), rabbit α-GAPDH (CST 14C10), rabbit α-LC3B (Novus NB100-2220), mouse α-EEA1 (BD Biosciences 610457), rabbit α-Golgin 97 (CST D8P2K), rabbit α-Citrate Synthase (CST D7V8B), rabbit α-GRP78 (Abcam ab21685), rabbit α-Histone H4 (CST D2X4V), rabbit α-Lamin A/C (CST 4C11), mouse α-ALIX (Santa Cruz Biotechnologies sc-53540), mouse α-Flotillin 2 (BD Biosciences 610384), α-CD63 (BD Biosciences, 556019), rabbit α-CD9 (CST D801A), rabbit α-ATG7 (CST D12B11), rabbit α-Phospho-S6K (CST 9205), rabbit α-S6K (CST 9202).

### EV isolation

EVs were isolated as described previously. Briefly, seven 150 mm culture dishes of HEK293T cells were cultured to 80% confluency in DMEM and 10% FBS, washed with 10 ml of PBS. Aliquots (30 ml) of EV depleted complete growth medium was added and EVs were collected after 48 h of cell growth or otherwise indicated. Conditioned medium was collected and subjected to sequential steps of differential centrifugation at 1000xg for 20 min, 10,000xg for 20 min and 100,000xg for 1 h. The pellet fractions collected after the final centrifugation was termed as high-speed pellet (HSP). To isolate CD63^+^ EVs, HSP was resuspended in 500 μl of PBS using gentle nutation for 1 h at 4°C on a rocking platform and incubated with 1 μg of monoclonal α-CD63 antibody (BD Biosciences) in presence of RiboLock (Thermo Scientific) RNase inhibitor at 1U/μL for one hour at 4°C. Aliquots (25 μl) of magnetic protein A/G beads (Pierce/Thermo Scientific) pre-equilibrated in PBS were added to the above reaction and further incubated for 1 h at 4°C. Beads were separated on a magnetic rack and washed three times with 500 μl PBS and resuspended in 100 μl of PBS, 20 μl of which was kept for immunoblotting and remainder was used for RNA analysis.

### Immunofluorescence

Immunofluorescence experiments were conducted as described before. Briefly, MDA-MB-231 cells were grown to about 30% confluency on poly-D-lysine coated coverslips. Cells were washed with PBS, fixed with 4% paraformaldehyde (Electron Microscopy Sciences) for 20 min at room temperature and then washed 3X with PBS. Following fixation, cells were permeabilized and blocked with a blocking buffer (PBS containing 2% FBS and 0.02% saponin) for 30 min at room temperature. Coverslips containing fixed and blocked cells were then moved into a humidity chamber and incubated with primary antibodies at 1:100 dilution in blocking buffer for 1 h at room temperature. Coverslips were washed three times with PBS and incubated with fluorophore-conjugated secondary antibodies at 1:500 dilution in blocking buffer for 1 h at room temperature. The coverslips were mounted onto glass slides using ProLong Diamond antifade reagent containing DAPI (Thermo Fisher Scientific) and sealed using a clear nail polish. LSM-900 microscope (Zeiss) equipped with a 63X Plan-Apochromat, NA 1.40 objective, was used to acquire images in the Airyscan 2 mode.

### OTTR Sequencing Pipeline

Since a large proportion of reads mapped to intergenic regions in the genome, all the reads were mapped to the curated and available transcriptome data. Although less likely based on our rigorous DNase treatment and lack of RT-PCR products in RT negative control lanes (Fig. 3), such reads could be a result of very low-level DNA contamination in our preparation. Another possibility is that OTTR-seq is able to pick up low-level background transcription at these intergenic loci.

Raw sequencing reads from OTTR libraries were trimmed and low-quality reads were removed using cutadapt:

cutadapt -q 10 -u 7 -a NNGATCGGAAGAGCACACG -m 15 -o path/to/output.fq.gz path/to/input.fq.gz

After creating a custom ncRNA library consisting of mature tRNA, mito-tRNA snoRNA, yRNA, scaRNA, rRNA, mito-rRNA, etc., an index was created using salmon:

salmon index -t ncRNA-library.fa -i ncRNA-index -k 19

Trimmed OTTR reads were aligned to the custom index using the salmon command,

salmon quant -i ncRNA-index -l SF -r /path/to/trimmedReads.fq.gz --validateMappings -o outputFolder

All scaRNA and snoRNA hits were validated by aligning trimmed OTTR reads to the genome using STAR as described below.

Because tRNAs and miRNAs may map poorly using traditional aligners, we specifically remapped these reads using tRAX and mirdeep2, respectively. After downloading mature tRNA files from GtRNAdb, a tRNA reference database was created with the following command: maketrnadb.py --databasename=/global/scratch/users/justin_krish/trax-genome/tRNAdb -- genomefile=hg38.fa --trnascanfile=hg38-tRNAs-confidence-set.out --gtrnafafile=hg38-mature- tRNAs.fa --namemap=hg38-tRNAs_name_map.txt

OTTR reads were mapped to tRNA sequences using the tRAX command, processsamples.py --experimentname=cytosolvsEndosome -- databasename=/path/to/users/tRNAdb --ensemblgtf=/path/to/hg38-genes.gtf -- samplefile=/path/to/samplefile.txt

For miRNA alignment, after downloading and extracting the human miRNAs from miRbase, miRNAs were aligned using the following commands,

mapper.pl /path/to/users/trimmedReads.fq -e -h -m -p /path/to/mirDeepIndex -s output.fa -t output.arf -v

miRDeep2.pl output.fa /path/to/genome.fa output.arf /path/to/mature-miRNAs.fa none

/path/to/hairpin-miRNAs.fa -t Human 2>1_report.log

### CORALL Sequencing Pipeline

Raw sequencing reads from OTTR libraries were trimmed and low-quality reads were removed using cutadapt:

cutadapt -q 10 -u 12 -a AGATCGGAAGAGC -A NNNNNNNNNNNNAGATCGGAAGAGC -o

/path/to/output/read1.fq.gz -p /path/to/output/read2.fq.gz /path/to/input/read1.fq.gz

/path/to/input/read2.fq.gz --minimum-length=1

Reads were mapped to the ncRNA transcriptome and quantified:

salmon quant -i /path/to/ ncRNA-index -l ISF -1 /path/to/read1.fq.gz -2 /path/to/read2.fq.gz -- validateMappings -o /path/to/output

Or to the genome using STAR:

STAR --runMode genomeGenerate --genomeDir /path/to/starGenomeOutput –

genomeFastaFiles /path/to/GenomeFiles/Homo_sapiens.GRCh38.dna_sm.primary_assembly.fa

--sjdbGTFfile /path/to/GenomeFiles/Homo_sapiens.GRCh38.103.gtf --sjdbOverhang 149

STAR --runMode alignReads --genomeDir /path/to/genomeIndex --readFilesIn

/path/to/trimmmedReads/read1.fq.gz /path/to/trimmmedReads/read2.fq.gz --outSAMtype BAM SortedByCoordinate --outFileNamePrefix /path/to/output --quantMode GeneCounts -- readFilesCommand zcat

Individual ncRNA hits were validated by comparing their alignment in Integrative Genomics Viewer (IGV) after creating an index of each bam file with samtools:

samtools index output.bam

## Acknowledgments

We dedicate this work to Bob Lesch, our lab manager for the past several decades who was tragically taken from us by an accident in 2021. We would like to thank all members of the Schekman lab for fruitful discussions during the preparation of this manuscript. We would also like to thank our current lab manager, Nam Che, Alison Killilea from the UC Berkeley Cell Culture Facility (SCR_017924) and Justin Choi from the UC Berkeley Functional Genomics Lab (SCR_022170). HEK293T-*atg7-/-* was a kind gift from the lab of Jayanta Debnath (Department of Pathology and Laboratory Medicine, UCSF). The mCherry-VPS4a (dominant mutant) cDNA was a gift from Kevin Rose in the laboratory of James H Hurley (UC Berkeley). The lab of K.C. is funded by the National Institutes of Health grant R35-GM127018 (E.N.), and the National Institutes of Health grant DP1 HL156819. R.S. is an Investigator of the Howard Hughes Medical Institute, a Senior Fellow of the UC Berkeley Miller Institute of Science and Chair of the Scientific Advisory Board of Aligning Science Across Parkinson’s Disease (ASAP). This work was funded by the Howard Hughes Medical Institute. The funders had no role in study design, data collection and interpretation, or the decision to submit the work for publication.

## Author Contributions

J.S., J.K.W., and R.S. designed the research; J.S., J.K.W., Q.E., and R.J. performed the research; L.F. and K.C. provided training and reagents; all authors analyzed the data; J.S., J.K.W., and R.S. wrote the manuscript, J.S., J.K.W., K.C., and R.S. edited the manuscript.

## Classification

Biological Sciences, Biochemistry.

