## Supplementary Figures for "Endosome transcriptomics reveal trafficking of Cajal bodies into multivesicular bodies"

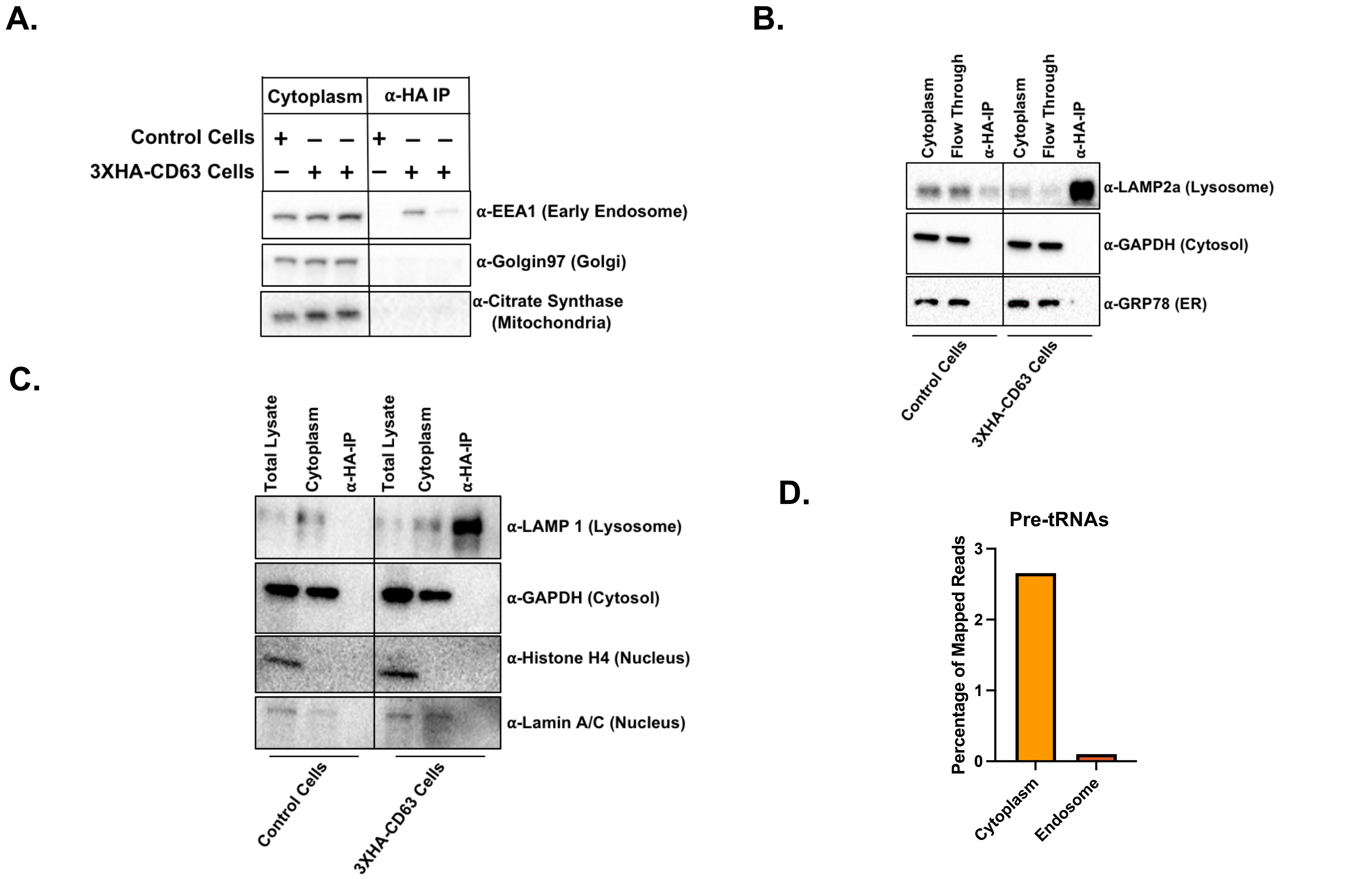
A.
Cytoplasm
α-HA IP
Control Cells
3XHA-CD63 Cells
α-EEA1 (Early Endosome)
α-Golgin97 (Golgi)
α-Citrate Synthase (Mitochondria)
B.
Cytoplasm
Flow Through
α-HA-IP
α-LAMP2a (Lysosome)
α-GAPDH (Cytosol)
α-GRP78 (ER)
Control Cells
3XHA-CD63 Cells
C.
Total Lysate
Cytoplasm
α-HA-IP
α-LAMP 1 (Lysosome)
α-GAPDH (Cytosol)
α-Histone H4 (Nucleus)
α-Lamin A/C (Nucleus)
Control Cells
3XHA-CD63 Cells
D.
Pre-tRNAs
Percentage of Mapped Reads
0
1
2
3
Cytoplasm
Endosome

Fig. S1

Figure S1. Control immunoblots and analysis of MVB quality. Immunoblots using markers for, A) Golgi, mitochondria and early endosomes; B) Lysosome, cytosol and endoplasmic reticulum; C) Lysosomes, cytosol and nuclei. D) A comparison of pre-tRNA levels in cytoplasm and MVBs/endosomes.

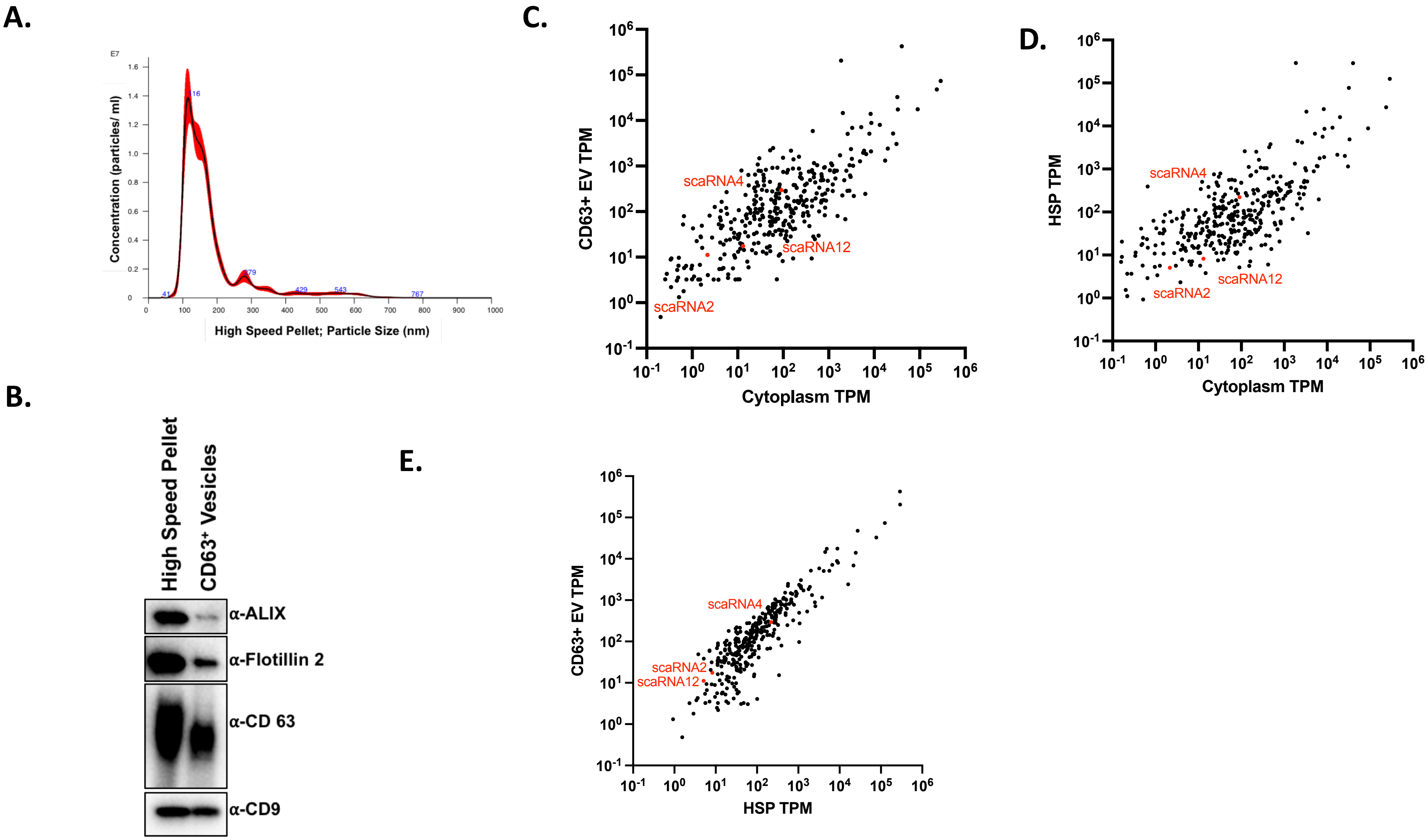
A.
E7
Concentration (particles/ ml)
High Speed Pellet; Particle Size (nm)
B.
High Speed Pellet
CD63+ Vesicles
α-ALIX
α-Flotillin 2
α-CD 63
α-CD9
C.
CD63+ EV TPM
Cytoplasm TPM
scaRNA4
scaRNA12
scaRNA2
D.
HSP TPM
Cytoplasm TPM
scaRNA4
scaRNA12
scaRNA2
E.
CD63+ EV TPM
HSP TPM
scaRNA4
scaRNA2
scaRNA12

Fig. S2

Figure S2. Purification and transcriptome of EVs. A) Nanoparticle-tracking of particles isolated by differential centrifugation of CM (HSP). B) Immunoblotting of HSP and $CD63^+$ extracellular vesicles (EV) for common EV markers. C) A comparison of transcriptome of cytoplasm to $CD63^+$ EVs and D) to the cytoplasm to HSP. E) Comparison of transcriptomes of $CD63^+$ EVs and HSP.

Fig. S3

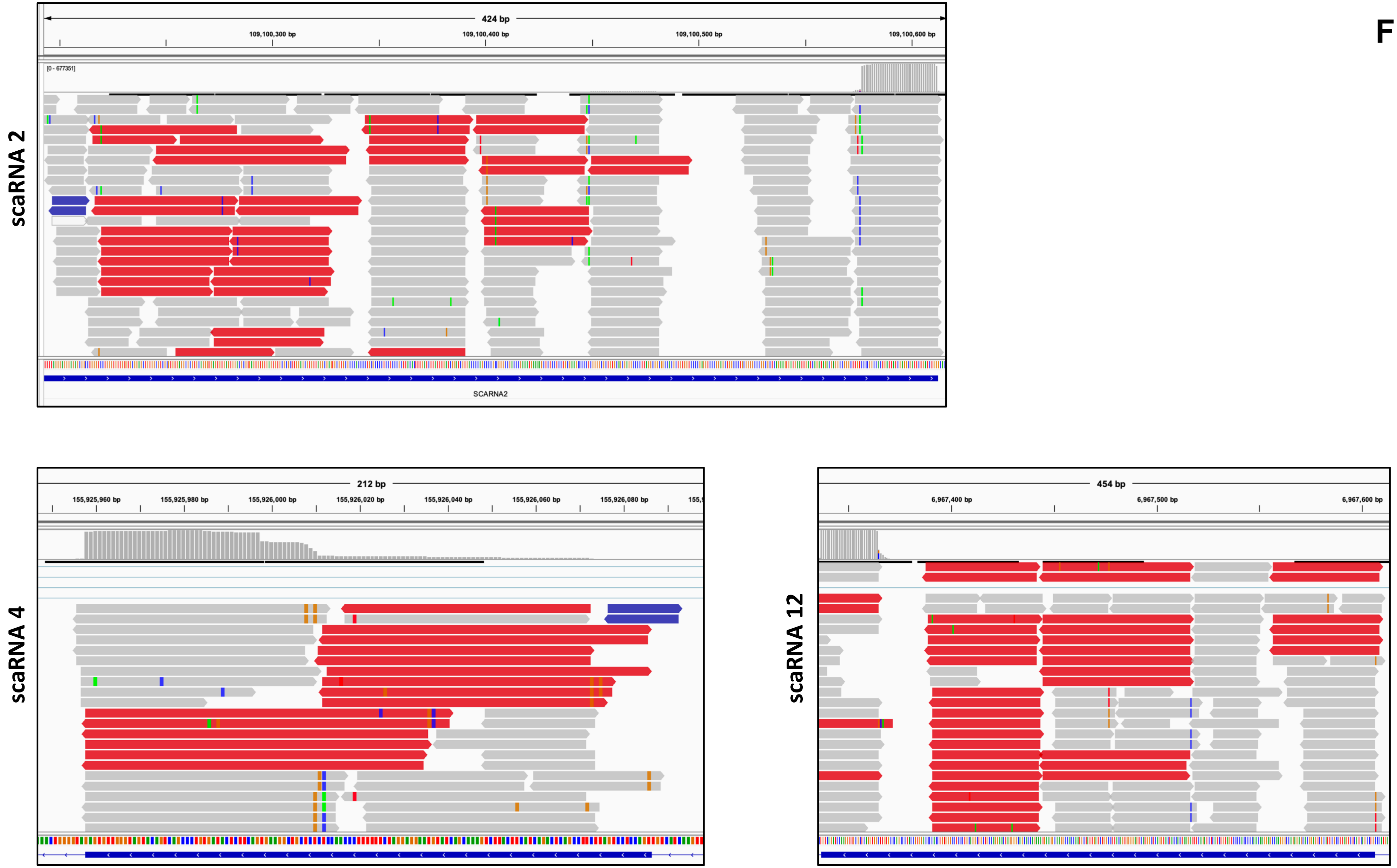
scaRNA 2
424 bp
109,100,300 bp
109,100,400 bp
109,100,500 bp
109,100,600 bp
[0 - 677351]
SCARNA2
scaRNA 4
212 bp
155,925,960 bp
155,925,980 bp
155,926,000 bp
155,926,020 bp
155,926,040 bp
155,926,060 bp
155,926,080 bp
scaRNA 12
454 bp
6,967,400 bp
6,967,500 bp
6,967,600 bp

Figure S3. Genome browser views of OTTR-seq reads for scaRNAs 2, 4 and 12.

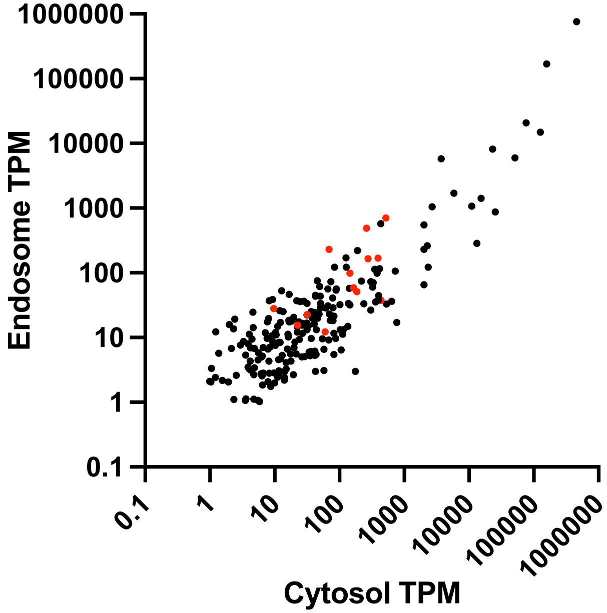
1000000
100000
10000
1000
100
10
1
0.1
Endosome TPM
0.1
1
10
100
1000
10000
100000
1000000
Cytosol TPM

**Fig. S4**

Figure S4. A comparison of conventional total RNA-seq reads obtained by CORALL-seq for MVB/endosome and cytoplasm samples. Highlighted in red are some of the snoRNAs and scaRNAs also detected using OTTR-seq.

**A.**

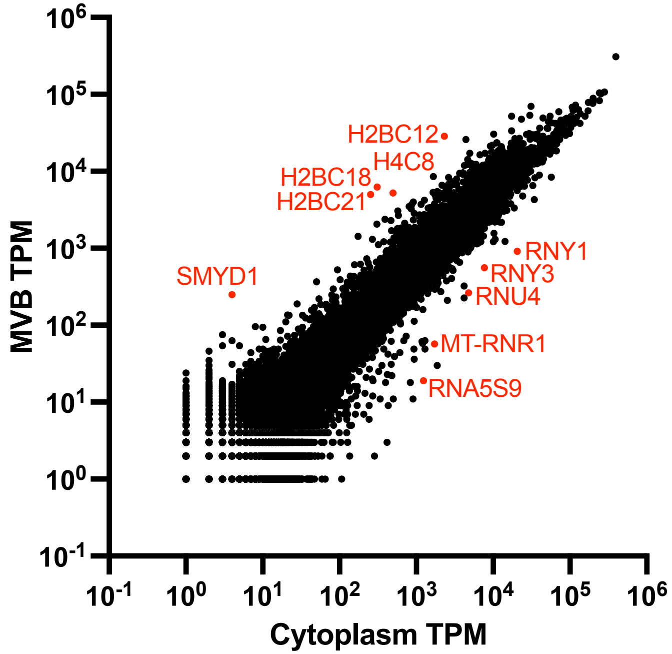


**B.**

| MVB-enriched gene | MVB-TPM |
|---|---|
| SMYD1 | 149.87 |
| HIST2H2BF | 76.28 |
| HIST2H2BE | 72.74 |
| HIST1H2BK | 55.01 |
| RPS4XP13 | 46.88 |
| HIST1H4H | 39.38 |
| AC091939.1 | 38.24 |
| RPS4XP21 | 33.13 |
| AC019117.1 | 31.18 |
| HIST2H2BA | 28.34 |
| AC084759.3 | 27.02 |
| HIST1H2BD | 25.85 |
| FTH1 | 25.68 |
| RPL36AP6 | 24.30 |
| BRINP1 | 23.79 |
| STK24 | 22.16 |
| AC022028.2 | 22.16 |
| CPSF3 | 20.44 |
| HIST1H2BO | 20.23 |
| HMGN2P19 | 19.27 |
| AC009102.1 | 19.27 |
| CRABP2 | 18.76 |
| LYZ | 18.71 |
| AC010327.4 | 18.55 |

**Fig. S5**

Figure S5. A) A comparison of mRNA content of MVBs and cytoplasm from CORALL-seq. B) A table of 25 most enriched transcripts in MVBs relative to cytoplasm as detected by CORALL-seq.

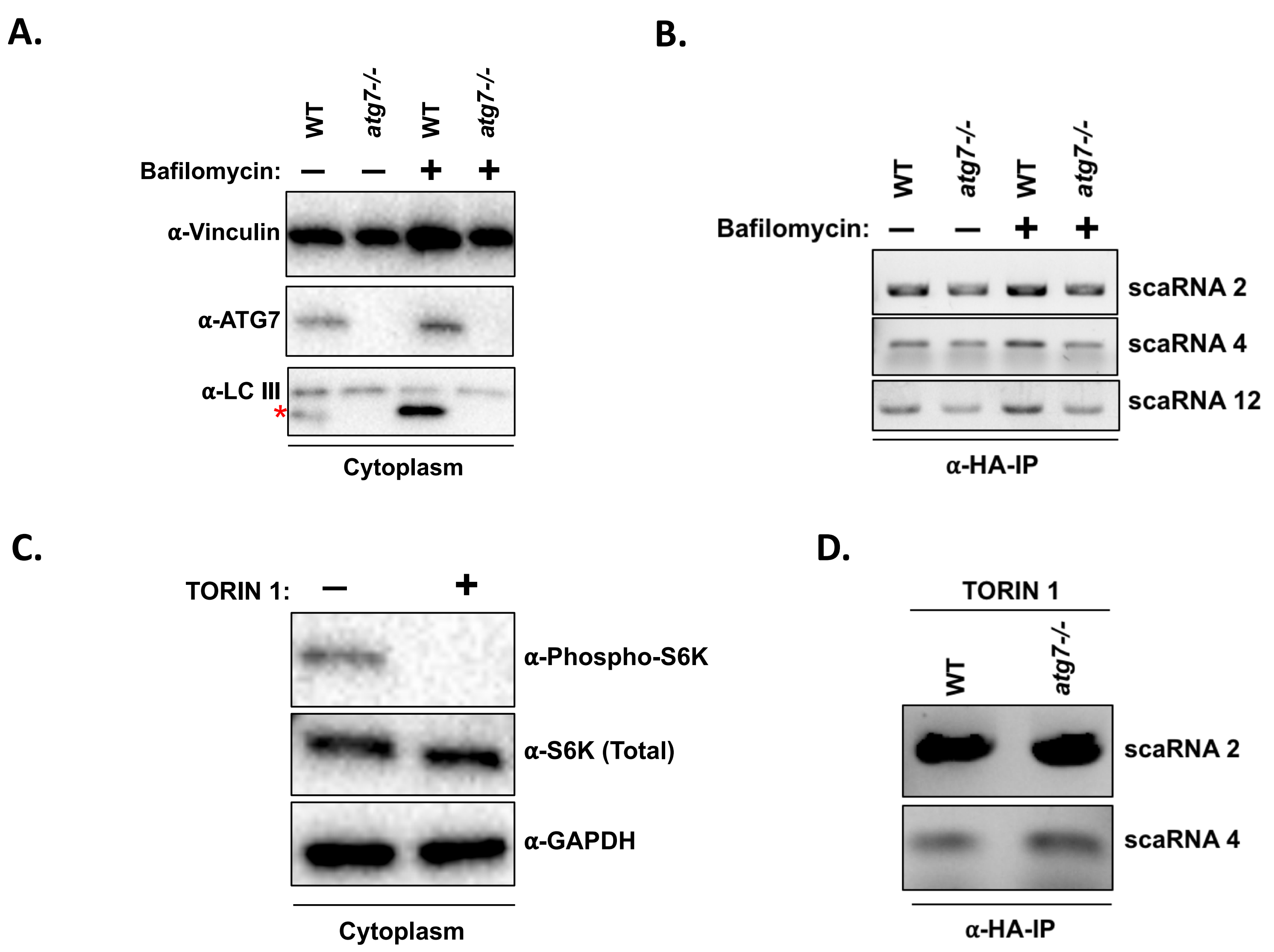
A.
WT
atg7-/-
WT
atg7-/-
Bafilomycin: — — + +
α-Vinculin
α-ATG7
α-LC III
*
Cytoplasm
B.
WT
atg7-/-
WT
atg7-/-
Bafilomycin: — — + +
scaRNA 2
scaRNA 4
scaRNA 12
α-HA-IP
C.
TORIN 1: — +
α-Phospho-S6K
α-S6K (Total)
α-GAPDH
Cytoplasm
D.
TORIN 1
WT
atg7-/-
scaRNA 2
scaRNA 4
α-HA-IP

Fig. S6

Figure S6. A) Immunoblots verifying atg7-/- allele and the resulting block of LC3 lipidation (band indicated by red asterisk. Bafilomycin treatments are as indicated. B) Analysis of scaRNA levels in MVBs by RT-PCR from treatments in A). C) Immunoblots verifying torin 1 treatment resulting in the block of S6K phosphorylation relative to total S6K levels. D) Analysis of scaRNA levels in MVBs by RT-PCR from treatments in C).
